## Supplementary Figures for "Functional networks of inhibitory neurons orchestrate synchrony in the hippocampus"

### Supporting information

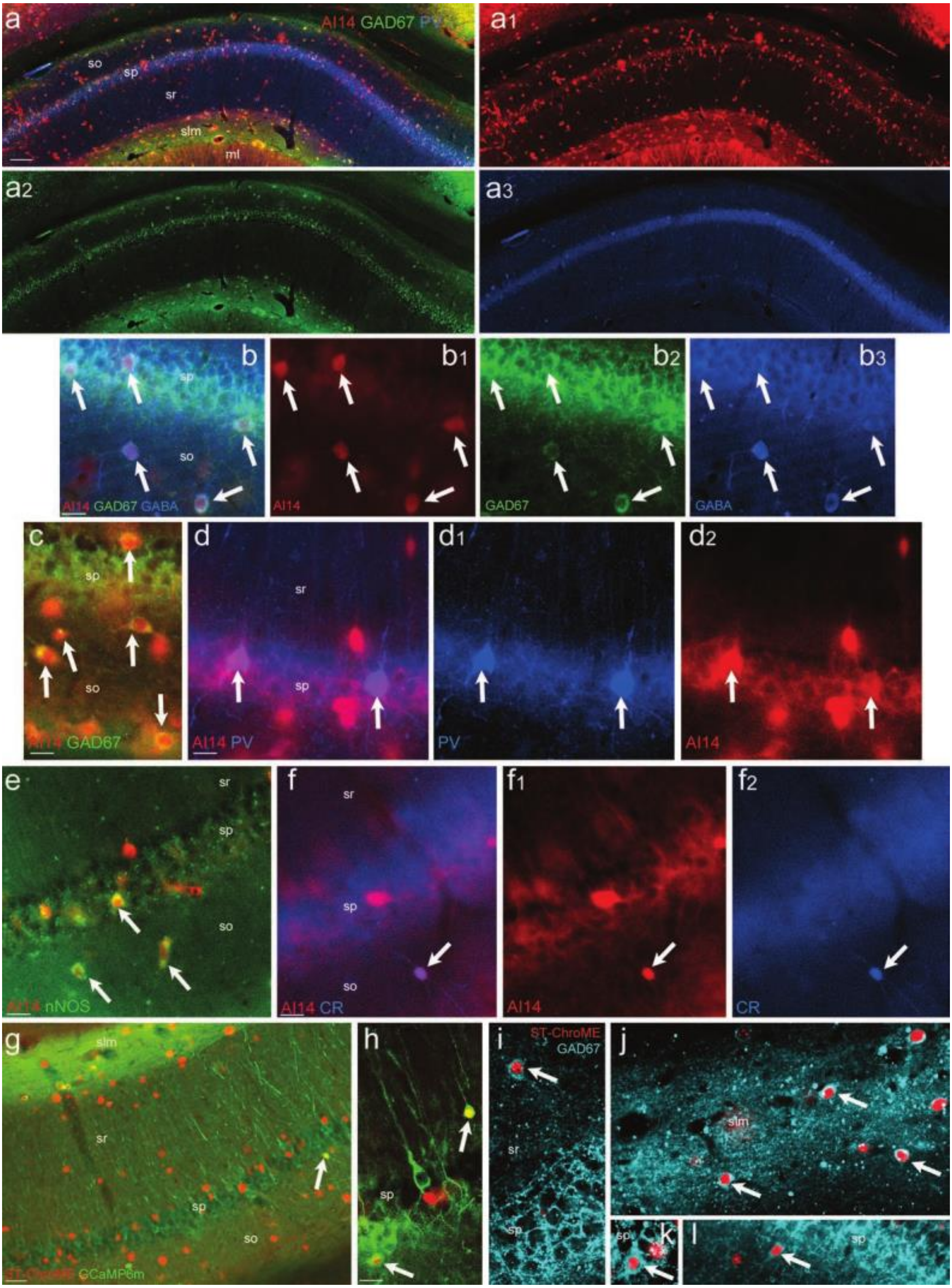

**Fig. S1** (related to Fig. 1, 2, 4) **- Validation of the GAD67-Cre transgenic mouse line to image CA1 interneurons.**

**a.** Td-Tomato (Ai14) neurons are distributed in all the CA1 layers and show the same pattern as GAD67-expressing cells. **b, c.** Ai14 cells are immunopositive (arrows) for GAD67 and GABA as shown in stratum pyramidale (sp) and stratum oriens (so). **d, e, f.** Some parvalbumin (PV), neuronal nitric oxide synthase (nNos), or calretinin (CR) cells are also expressing Ai14 (arrows in d, e, f, respectively). **g, h.** injection of cre-dependent ST-ChroME virus induces labeling of GABA neurons distributed as expected in all the layers of CA1, injection of GCaMP6m induces green fluorescent protein expression in both pyramidal neurons and some ST-ChroME positive cells (arrows). **i-l.** GAD67 immunolabeling confirms that ST-ChroME-positive cells are GABAergic neurons (arrows). slm, stratum lacunosum-moleculare; sr, stratum radiatum, ml, stratum moleculare. Scale bars, a, g: 100 µm; b, e, f, h-l: 20 µm.

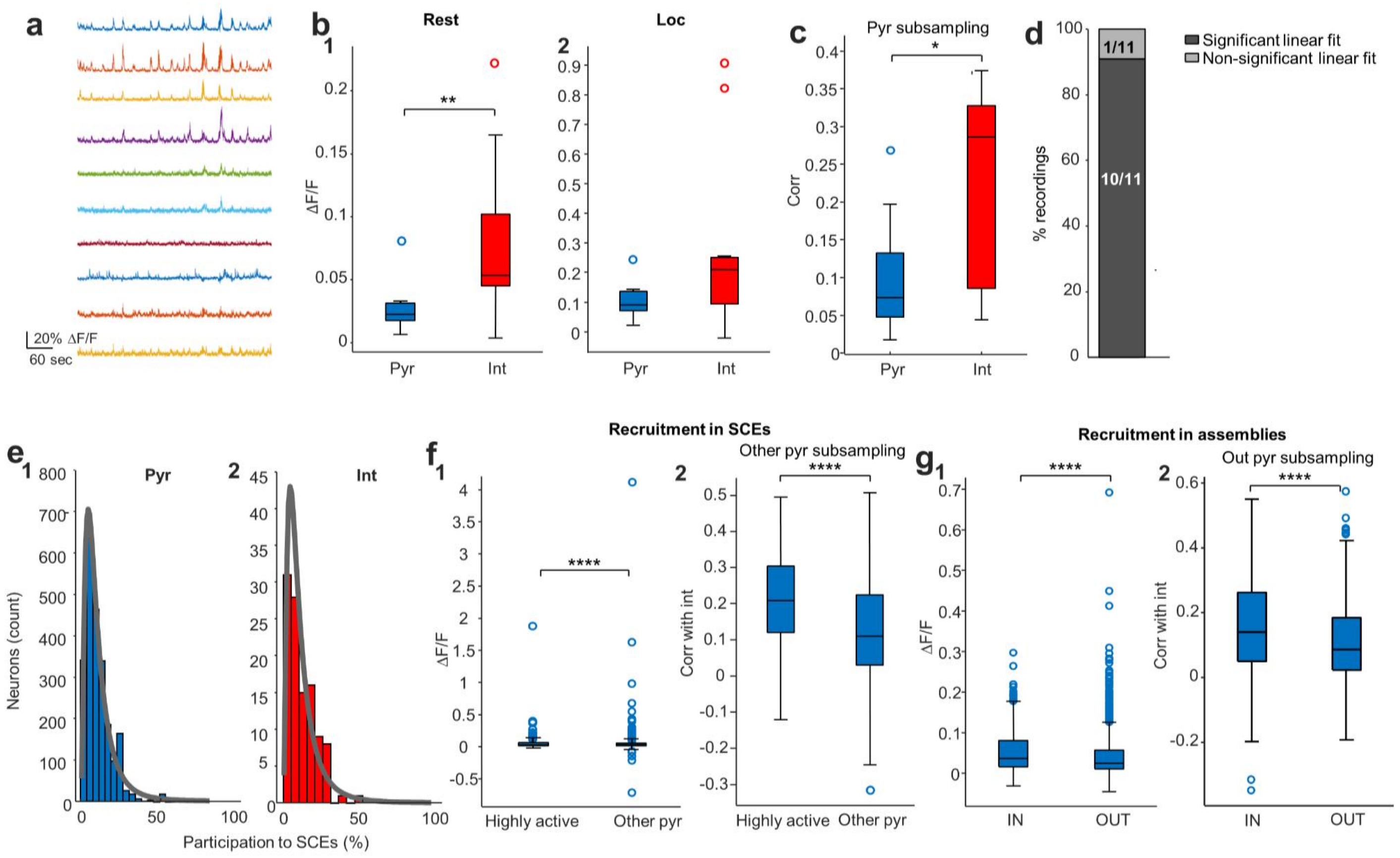


**Fig. S2** (related to Fig. 1) **- Activity of pyramidal cells and interneurons in relation to locomotion and SCEs.**

**a.** Calcium traces from ~7 min of recording from all interneurons in a representative field of view (10 interneurons in total). **b.** Same as Fig. 1c, but restricted to rest or locomotion periods. Interneurons show higher activity than pyramidal cells during rest periods (b1, p = 0.009), but not locomotion periods (b2, p = 0.115, both Wilcoxon signed rank tests, n=11 FOVs from 6 mice). **c.** Pairwise correlations between interneurons are significantly higher than the ones between pyramidal neurons (p = 0.041, Wilcoxon signed rank test, n=11 FOVs from 6 mice) even when subsampling pyramidal cells to match interneurons' ΔF/Fs (to control for the higher ΔF/F of interneurons). **d.** Linear model fitted for pyramidal-pyramidal vs pyramidal-interneurons pairwise correlations (see Fig. 1h) for individual recordings: proportion of recordings displaying significant fit (p<0.05). **e.** Distribution of the proportion of SCEs to which each cell participates. Both pyramidal cell (d1) and interneuron (d2) distribution show lognormal shapes. Lognormal fits are depicted in grey. **f1.** Pyramidal cells that are highly active in SCEs (scoring above the 90th percentile in the distribution of SCE participation including all pyramidal cells) display significantly higher ΔF/F than other pyramidal cells (p = 0.008, Mann-Whitney U test, n=276 highly active cells, n=2517 other cells, from 11 FOVs from 6 mice). **f2.** Pyramidal cells that are highly active in SCEs have significantly higher pairwise Pearson's correlations to interneurons compared to other cells even when other cells are subsampled to match highly active cells' ΔF/Fs ($p=9.7e^{-14}$, Mann-Whitney U test, n=276 highly active cells, n=275 other cells, from 11 FOVs from 6 mice). **g1.** Pyramidal cells that are part of cell assemblies (IN) display significantly higher ΔF/F than pyramidal cells not in assemblies (OUT, $p = 4.2e^{-6}$, Mann-Whitney U test, n=361 in assemblies, n=1437 not in assemblies, from 11 FOVs from 6 mice). **g2.** Pyramidal cells that are part of cell assemblies (IN) display significantly higher ΔF/F than pyramidal cells not in assemblies (OUT, $p = 5.6e^{-6}$, Mann-Whitney U test, n=361 in assemblies, n=319 not in assemblies, from 11 FOVs from 6 mice). * p < 0.05; ** p < 0.01. Boxplots represent medians (center) and interquartile ranges (bounds). The whiskers extend to the most extreme data points not considered outliers, which are plotted individually using the circles. Underlying data can be found in S5 data.

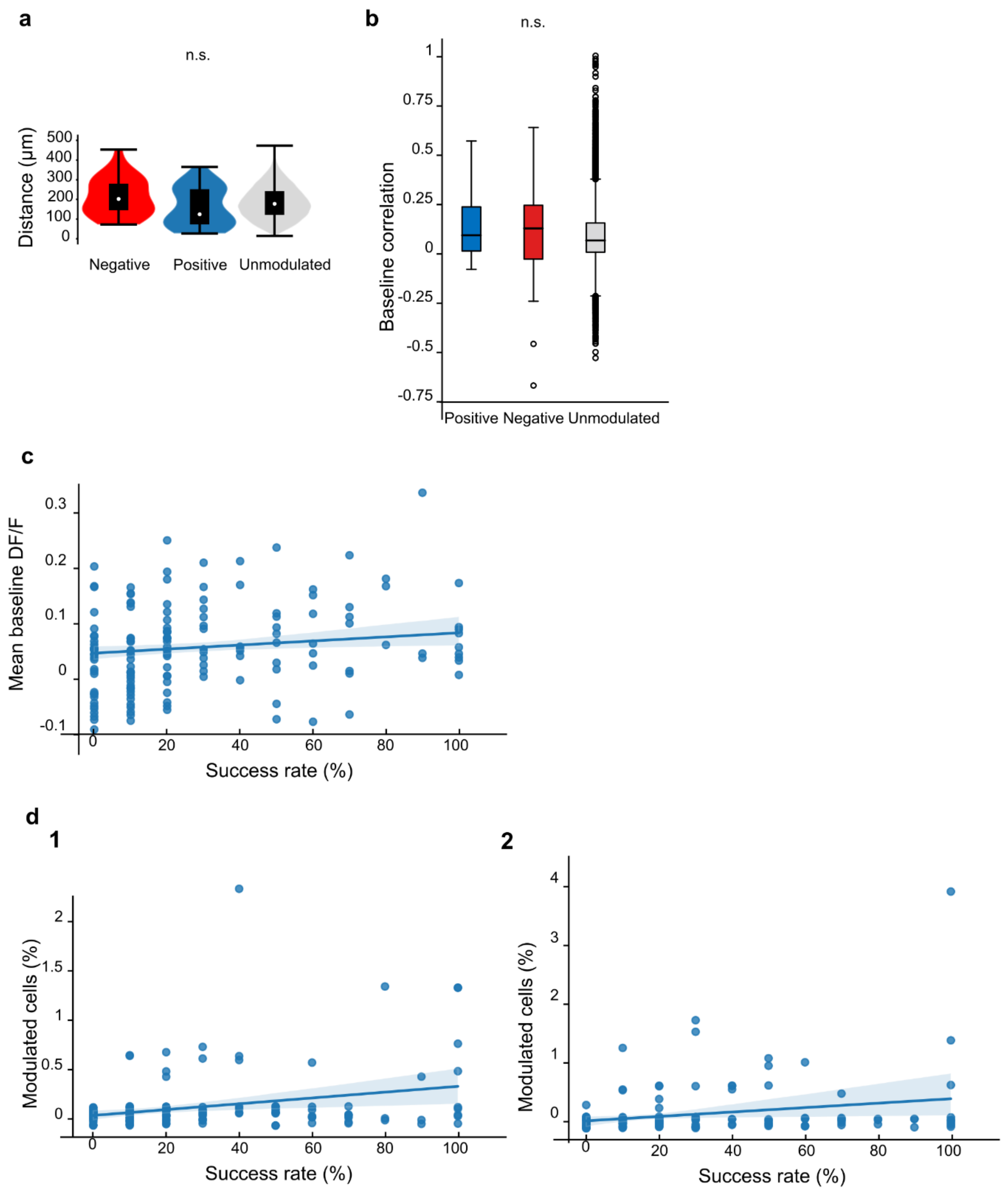


**Fig. S3** (related to Fig. 2) **- Indirectly modulated cells: spatial distribution, correlation, and link to success rate of the stimulated neuron**

**a.** Distribution of the distance between the stimulated interneuron and negatively (red), positively (blue) or unmodulated (gray) cells (Kruskal–Wallis H-test, three groups, p=0.19). **b.** Box plot indicates the correlation between the fluorescence calcium traces of positively (blue), negatively (red), and unmodulated (gray) neurons (Kruskal–Wallis H-test, three groups, p=0.17). **c.** Scatter plot with linear regression best-fit line indicating a significant correlation between the stimulation success rate and the mean baseline dF/F signal of the stimulated neuron (Pearson r=0.206, p=0.012) **d.** Scatter plot with linear regression best-fit line indicating a correlation between the fraction of positively (**d1**) or negatively (**d2**) modulated neurons and the success rate of the target cell (Pearson's r=0.251, p=0.002; Pearson's r=0.295, p=0.0003, respectively). Boxplots represent median (center) and interquartile ranges (bounds). The whiskers extend to the most extreme data points not considered outliers, which are plotted individually using the circles. Underlying data can be found in S6 data.

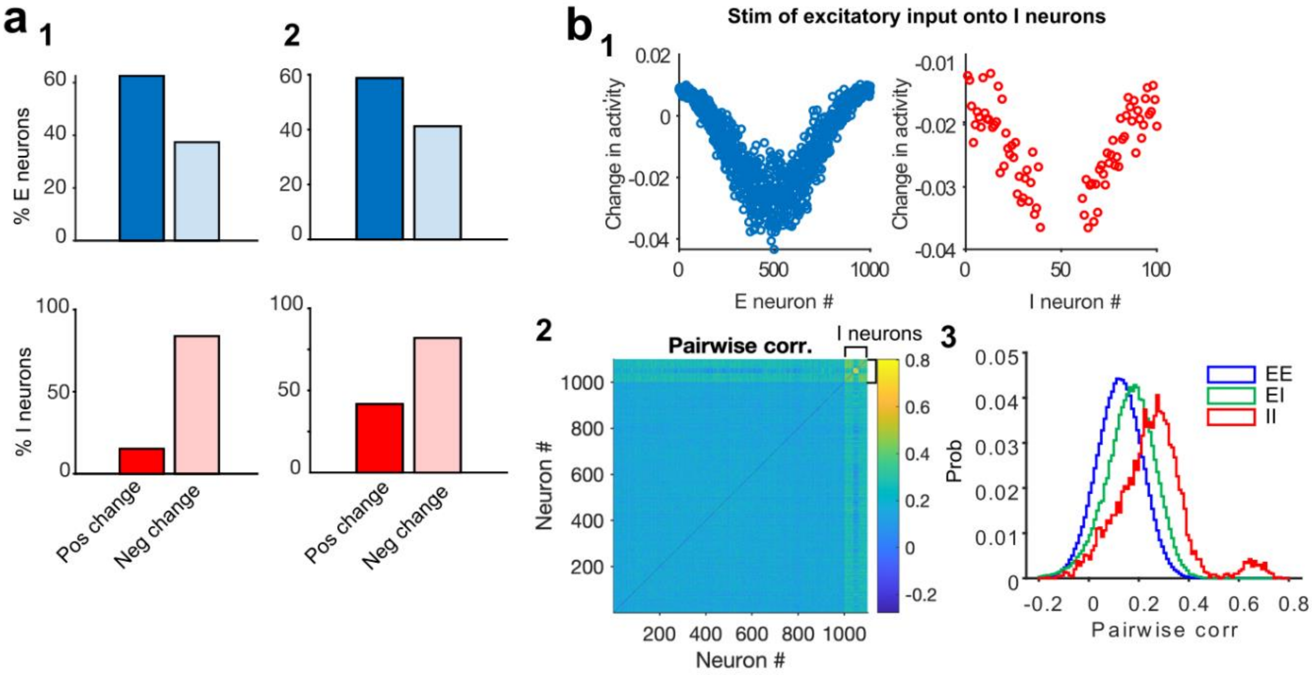


**Fig. S4** (related to Fig. 3) **- Further characterization of connectivity in the network model**

**a.** Fractions of E and I neurons showing a net positive or negative change in their activity, as a result of single I perturbations. **a1**.Results obtained when the effect is assessed from a linear analysis of the network dynamics and its weight matrix (W). **a2.** Results obtained when I-I connections display the same specificity as E-I connections (m_II = 1). **b.** Same as Fig. 3d, but with external stimulation of inhibitory neurons (20 I neurons in the middle; the changes in the activity of stimulated I neurons are not shown). Underlying data can be found in S7 data.

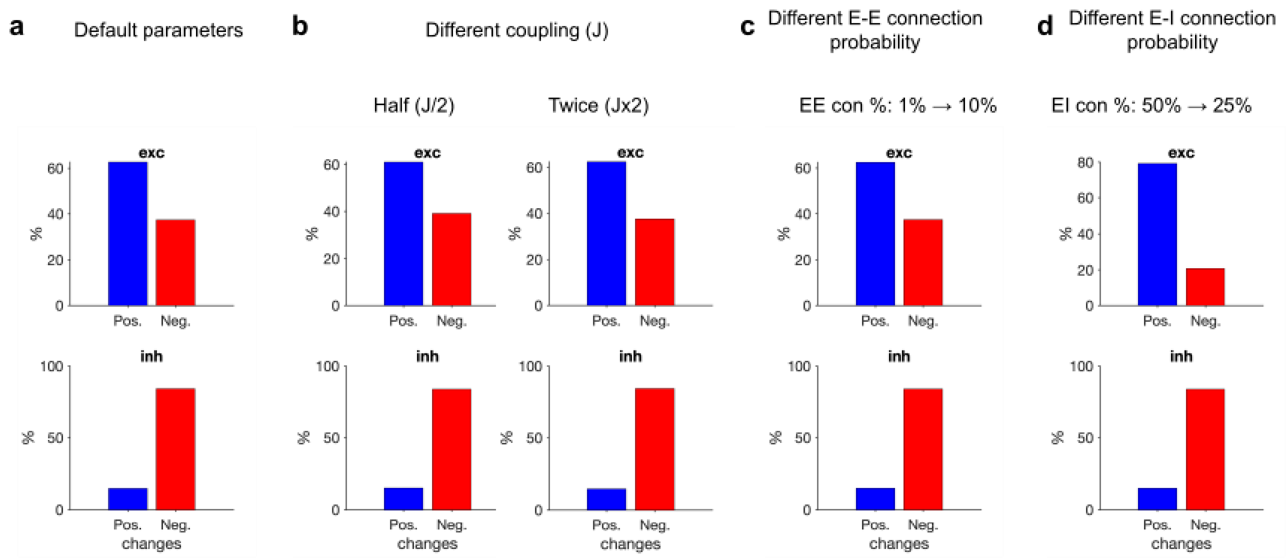


**Fig. S5** (related to Fig. 3) **- Robustness of the modelling results to network parameters.**

The results of our single inhibitory neuron perturbations were robust to the choice of network parameters, and fine-tuning was not needed to obtain the key results. To show the robustness of our results to the change of parameters, we simulated our networks with different ranges of parameters and calculated the modulations in each network. **b.** We decreased the main coupling in the network (J) by half or increased it twice, and observed similar results. **c**. We also changed the connection probability of E-E and E-I connections. We increased the connection probability of initially sparse E-E connections, from 1% to 10%, and observed similar results. d. We also decreased the connection probability of E-I connections, from the original 50% to 25%, and observed similar results. We therefore conclude that our results are robust to the choice of parameters in the network as our results hold for a wide range of parameter space. Underlying data can be found in S8 data.

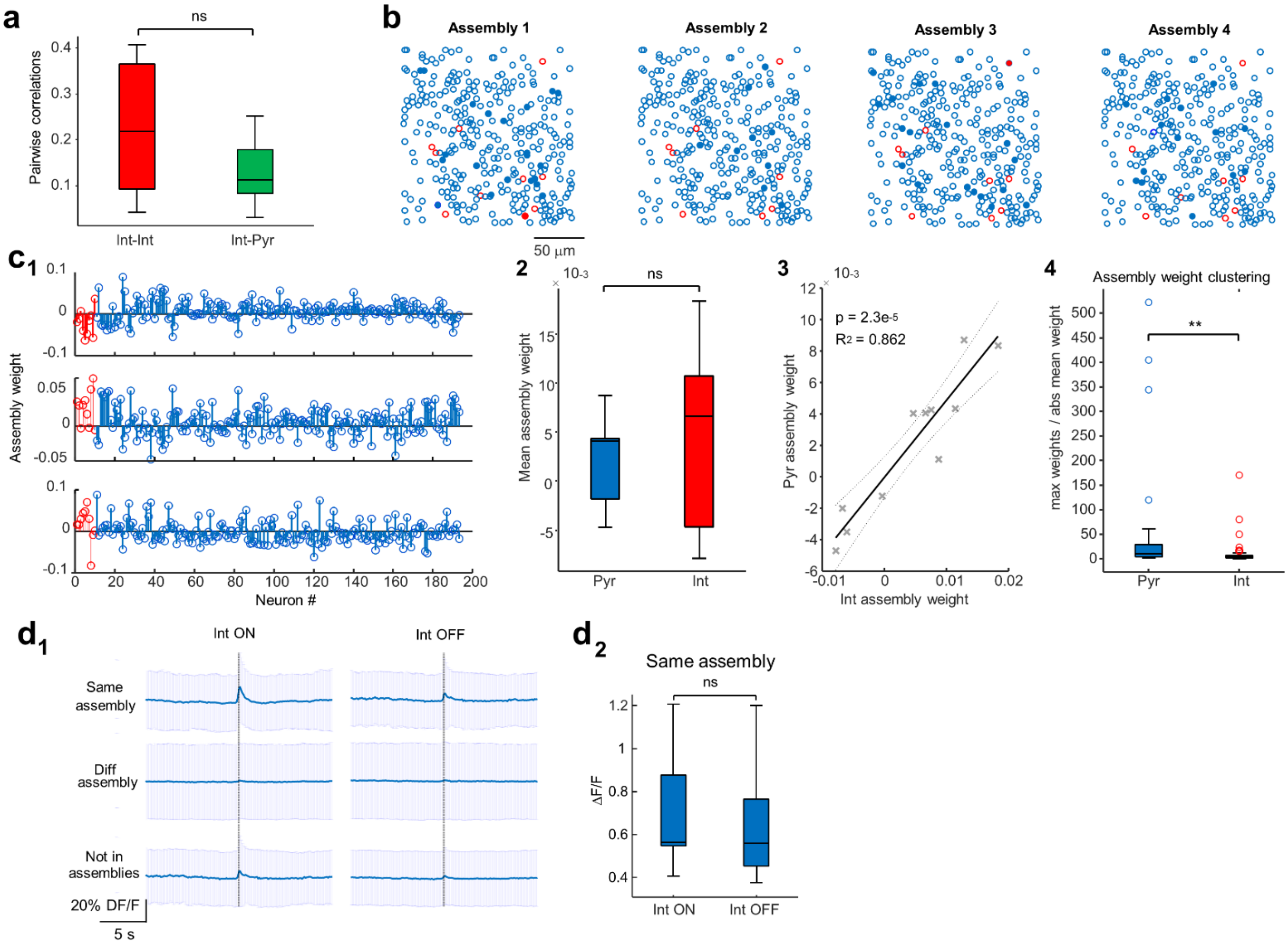


**Fig. S6** (related to Fig. 4) **- Relationship between interneuron activity and cell assemblies**

**a.** Interneurons are not clustered into single assemblies, as evidenced by the fact that pairwise correlations between interneurons are not higher than correlations between interneurons and pyramidal cells (p = 0.16791, Wilcoxon signed rank tests, n=11 FOVs from 6 mice). **b.** Contour maps indicating the centroids of all active neurons in a representative imaging session with four cell assemblies (SCE-based method). Pyramidal cells are depicted in blue, interneurons in red. Filled contours belong to an individual assembly, whereas empty contours do not. Note the lack of spatial clustering: pyramidal cells and interneurons, as well as cells forming and not forming assemblies, are intermingled. **c1.** Assembly patterns (from 3 significant cell assemblies) were obtained from a representative recording (same imaging session as c1) using the PCA/ICA assembly detection method (see Methods for details). Each plot represents a significant principal component (assembly), with a given weight for each neuron. Interneurons' weights are depicted in red, pyramidal cells' weights in blue. **c2.** Pyramidal cells and interneurons display similar assembly weights (p=0.4, Wilcoxon signed rank test, n = 31 assemblies from 11 recordings from 6 mice). **c3.** Fit of linear model between pyramidal cell assembly weight and interneuron assembly weight for each recording (averaged across 31 assemblies, n = 11 recordings, 6 mice). **c4.** Pyramidal cell assembly weights are more clustered in a single assembly compared to interneurons (maximum assembly weight across assemblies divided by the absolute average of assembly weights; p = 0.0025, Mann Whitney U test, n = 31 assemblies, from 11 FOVs from 6 mice). **d1.** Lack of evidence of cell assembly segregation by single interneurons. Assembly activation-triggered average of pyramidal cells' calcium traces when each interneuron in an assembly is active (left) or inactive (right). Shaded areas represent standard deviations. *Top*, traces from pyramidal cells in the same assembly as the interneuron. *Middle*, traces from pyramidal cells in different assemblies. *Bottom*, traces from pyramidal cells not forming assemblies. Note that the activity of the interneuron in an assembly does not affect the activity of pyramidal cells of competing assemblies or of the ones not forming assemblies. **d2.** No significant difference in ΔF/F peak at assembly activation for pyramidal cells when the interneuron in the same assembly is active or inactive ($p=0.7$, Wilcoxon signed-rank test, $n=7$ recordings with significant assemblies from 5 mice). Data from 21 cell assemblies. ** $p < 0.01$. Boxplots represent medians (center) and interquartile ranges (bounds). The whiskers extend to the most extreme data points not considered outliers, which are plotted individually using the circles. Underlying data can be found in S9 data.

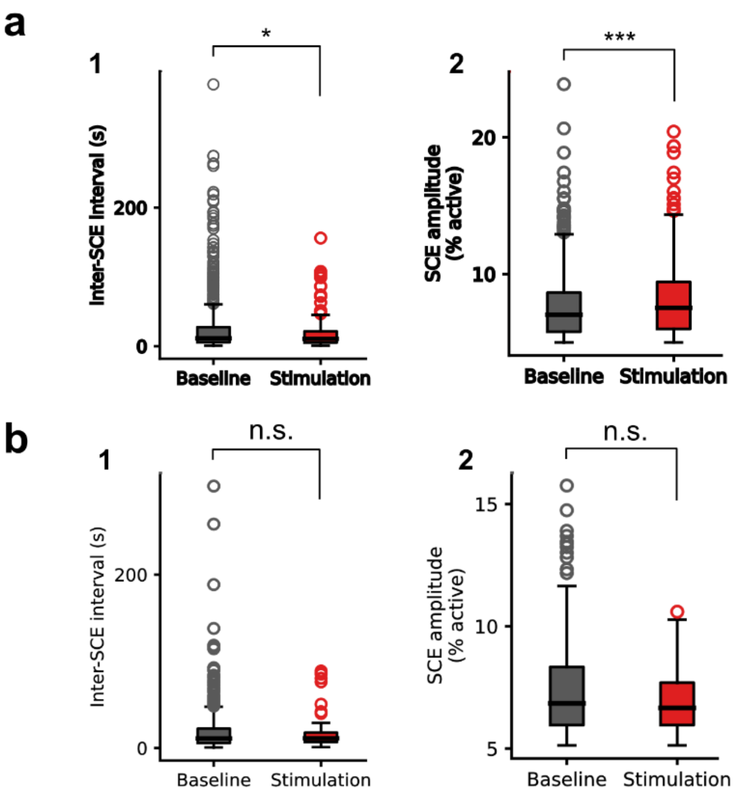


**Fig. S7** (related to Fig. 4) **- Changes in SCE frequency and amplitude are absent if targeted interneuron does not respond to light stimulation**

**a1.** Same as Fig. 4b3, but for all the experiments, excluding the experiments with unresponsive cells, p=0.035. **a2.** Same as Fig. 4 b4, but for all the experiments, excluding the experiments with unresponsive cells, p=0.0007. **b1.** Same as Fig. 4b3, but for the experiments with unresponsive cells only, p=0.45. **b2.** Same as Fig. 4 b4, but for the experiments with unresponsive cells only, p=0.13 (Mann Whitney U test in all cases). * $p < 0.05$; *** $p < 0.001$. Boxplots represent medians (center) and interquartile ranges (bounds). The whiskers extend to the most extreme data points not considered outliers, which are plotted individually using the circles. Underlying data can be found in S10 data.

**Supplementary table.**

Table of the distribution of all-optical experiments and modulated cells per animal.

| | | | | Direct response | Indirect responses | |
|---|---|---|---|---|---|---|
| Mouse ID | n FOVs | n stimulated interneurons | Total n Int | n stimulation trials with response | Total n positively modulated neurons | Total n negatively modulated neurons |
| **ID_1** | 11 | 39 | 267 | 119 | 22 | 4 |
| **ID_2** | 9 | 32 | 193 | 61 | 2 | 3 |
| **ID_3** | 6 | 22 | 126 | 74 | 1 | 4 |
| **ID_4** | 8 | 18 | 107 | 101 | 4 | 11 |
| **ID_5** | 8 | 15 | 76 | 21 | 1 | 0 |
| **ID_6** | 4 | 10 | 47 | 41 | 9 | 1 |
| **ID_7** | 2 | 4 | 17 | 7 | 0 | 0 |
| **ID_8** | 1 | 3 | 26 | 4 | 0 | 0 |
| **ID_9** | 2 | 3 | 10 | 1 | 0 | 0 |
| **ID_10** | 1 | 2 | 6 | 3 | 0 | 0 |
| **ID_11** | 1 | 1 | 4 | 1 | 0 | 0 |
